# Model-based control of individual finger movements for prosthetic hand function

**DOI:** 10.1101/629246

**Authors:** Dimitra Blana, Antonie J. van den Bogert, Wendy M. Murray, Amartya Ganguly, Agamemnon Krasoulis, Kianoush Nazarpour, Edward K. Chadwick

**Affiliations:** Institute for Science and Technology in Medicine, Keele University, UK; Cleveland State University, USA; Northwestern University and the Shirley Ryan Ability Lab, USA; Newcastle University, UK

## Abstract

Prosthetic devices for hand difference have advanced considerably in recent years, to the point where the mechanical dexterity of a state-of-the-art prosthetic hand approaches that of the natural hand. Control options for users, however, have not kept pace, meaning that the new devices are not used to their full potential. Promising developments in control technology reported in the literature have met with limited commercial and clinical success. We have previously described a biomechanical model of the hand that could be used for prosthesis control. In this study, we report on three key elements of the biomechanical simulations relevant to prosthesis control: we show the performance of the model in replicating recorded hand kinematics and find average correlations of 0.89 between modelled and recorded motions; we show that the computational performance of the simulations is fast enough to achieve real-time control with a robotic hand in the loop; and we describe the use of the model for controlling object gripping. Despite some limitations in accessing sufficient driving signals, the model performance shows promise as a controller for prosthetic hands when driven with recorded EMG signals. We identify areas for future work to address these limitations.

## I. Introduction

**D**EXTEROUS and natural finger movement, including manipulation of objects, is an important goal for upper limb prosthesis users [1]. Increasingly sophisticated prosthetic devices have become available over the last few years, offering individual digit movement and degrees of freedom approaching those of the natural hand [2]. However, a lack of sophisticated control options for users limits full exploitation of these devices, and control is characterized by predefined patterns of grasp and sequential actions [3].

With access to more input signals from muscles [4] or nerves [5], the potential for natural and simultaneous control of multiple degree-of-freedom (DOF) movement is increasing. However, for this to become a reality, an intuitive means of control for these sophisticated devices is needed. We have proposed the use of a biomechanical hand model as a controller for the prosthetic device whereby control signals based on electromyography (EMG) recorded from the user’s residual muscles drive a dynamic simulation of hand motion [6]. The resulting modelled digit actions, based on the biomechanics of the natural hand, can be replicated in real time by the prosthesis, producing natural hand movements.

The current state-of-the-art in myoelectric prosthesis control is dominated by machine learning techniques, whereby a decoding algorithm maps residual muscle signals to desired actions. This approach presupposes no particular relationship between the muscle signals and desired actions, but trains the controller by recordings made from prosthesis users attempting to carry out desired actions [7]. Training on data recorded during dynamic movements, rather than being limited to static postures, has been shown to improve the robustness of these systems [8], [9] and users have been shown to adapt to the dyanamics of a physcial device as well as controlling kinematic signals [10]. In addition to mapping different grip postures, recent work has also attempted to map surface EMG signals to individual finger movement using various pattern recognition algorithms [4], [11], [12]. In some cases, where access to the recording sites is lost as a consequence of the amputation, targeted muscle reinnervation may be used to transfer residual nerves into alternative muscles. These can then be used as recording sites for EMG sensors to produce the control command [13].

Some commercial systems are available using this type of technology, however its use requires significant training on the part of the user. More commonly seen in clinical or commercially available devices is proportional control, where a user can modulate the degree of movement (speed, angle, force) by controlling the amplitude of generated muscle signals [14]. In order to achieve multiple grip patterns, these systems require mode switching between grips, or sequential control of single degrees of freedom.

In our approach, we take advantage of the known biomechanics of the limb to simulate the actions that would result from particular muscle activation patterns if the limb were still present. As long as the biomechanical model is a reasonable approximation of the missing limb, the EMG-driven, simulated movements should be a good approximation of the desired movements. Using the biomechanics of the limb in this way to interpret residual muscle signals and generate movement commands may help to reduce the uncertaintly associated with noisy measurements of muscle activation and reduce the ambiguity of intended actions. A further benefit of this approach is that the explicit representation of muscle elements in the model, in contrast to pure machine learning approaches, allows the generation of proprioceptive signals that can be fed back to the prosthesis user to provide truly closed-loop control of movement [15]. Muscle length and velocity information provided by modelled muscle spindle output, and tendon force feedback generated by a simple model of the Golgi Tendon Organ (GTO), could be given to the user via peripheral nerve stimulation.

Several recent studies have shown the potential for this model-based approach in controlling wrist and hand movement from recorded EMG signals. Crouch & Huang [16] used a real-time, two-DOF musculoskeletal model to control the fingertip of a virtual hand with EMG signals from four forearm muscles. Sartori *et al.* [17] included an EMG-driven musculoskeletal model that decoded joint moments in a control scheme for wrist movement and hand opening-closing and Kapelner *et al.* [18] have attempted to improve the human-machine interface by decomposing recorded EMG signals into the underlying neural drive. This neuromechanical approach shows promise for real-world applications due to the robustness of the control to movement artifacts and the physiologically-constrained solution space for control signals.

We have previously shown stand-alone biomechanical hand model simulations of individual finger movements that run faster than real time [6]. In this study, we attempt to answer three key questions that will enable translation of the theoretical model approach to actual device implementation:

1. Are simulated model movements the same as those intended by the user?
2. Can a model of realistic complexity, with hardware in the loop, run fast enough to enable real-time control?
3. For interaction with the environment, could the model be used to control object gripping as well as open-chain movements?

To answer these three questions, we describe this study in three parts: part 1 compares the kinematics of natural hand movement against those of an EMG-driven biomechanical model in a group of normally-limbed individuals; part 2 drives a robotic hand with the EMG-controlled biomechanical model, and assesses the model in terms of its computational speed; and part 3 demonstrates the use of the model in simulated gripping of a cup being filled with liquid.

## II. Methods

### A. Biomechanical hand model

The biomechanical hand model is a modified version of the one we have presented previously [6]. OpenSim [19] is used to modify and visualise the structure of the model; the metacar-pophalangeal (MCP) joints are modelled as two orthogonal hinge joints, the proximal and distal interphalangeal (PIP, DIP) joints are simple hinges, and the muscle lines of action are represented by elements passing from origin to insertion via wrapping objects to achieve the correct moment arms for the muscles across a range of postures. The multibody dynamics are described by the equation of motion:

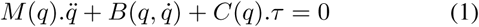

where *M* is the mass matrix; the 2^*nd*^ term accounts for centrifugal, Coriolis and gravitational forces and the final term includes the effects of joint moments *τ* via the coefficient matrix *C. τ* is the summation of muscle moments and passive joint moments. To speed up the computational performance of the model, the muscle lengths and lines of action are preprocessed to a polynomial representation to avoid the run-time calculation of wrapping paths [20].

To match the structure of the robotic hand used as part of the study (Section II-C), the model wrist was fixed, all finger abduction/adduction degrees of freedom were removed, and all muscles crossing only the wrist were removed. This simplifies the model somewhat compared to the full model published in Blana *et al.* [6], and makes control with surface EMG recordings more feasible. The resulting model has 16 degrees of freedom (four at the thumb, three at each of the fingers) and 18 muscles (see Fig. 1).

**Fig. 1.**
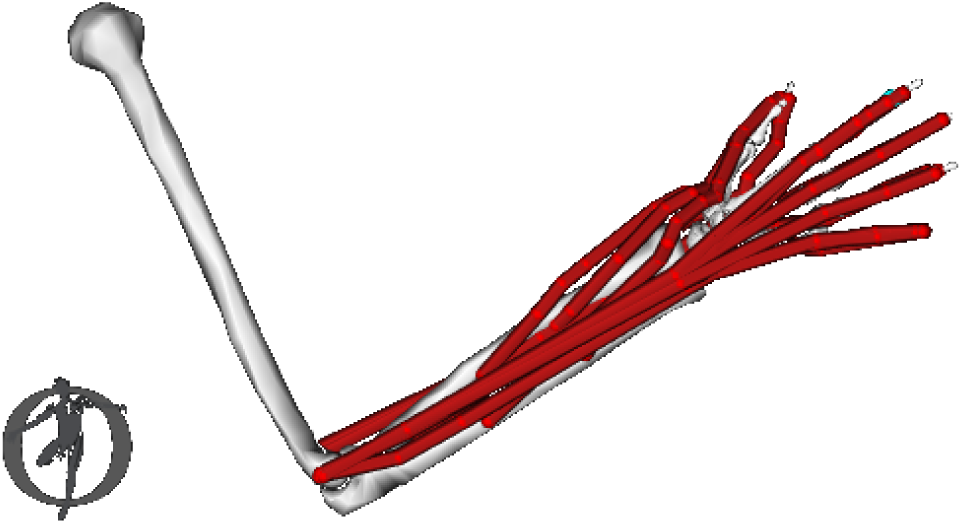
OpenSim visualisation of the hand model, showing muscle lines of action and included joints (CMC, MCP, IP at the thumb, MCP, PIP and DIP for the fingers).

Muscle dynamics are simulated with a first order delay, and integration of system dynamics is carried out using an implicit formulation to allow for larger stable integration step sizes:

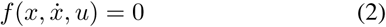

The state vector *x* contains 68 variables: 16 angles *q*, 16 angular velocities 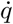, 18 muscle contraction state variables *s*, and 18 muscle active states *a*. The implicit formulation of system dynamics enables faster than real time, forward-dynamic simulation but requires explicit calculation of Jacobians 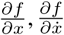 and 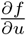. These were implemented in Autolev and hand-coded for muscle dynamics [21]. The model was implemented in OpenSim, and real-time simulations were run in Matlab (Mathworks, Inc., Natick, MA).

To simulate proprioceptive feedback signals, we added the computationally efficient models of proprioception developed by Williams & Constandinou [15]. The muscle spindle model is based on Mileusnic *et al.* [22] and is composed of three intrafusal fibre models (bag1, bag2 and chain) and two afferent firing models, primary (Ia) and secondary (II). This generates an output signal modulated by both muscle length and velocity. The Golgi tendon organ is modelled as a force sensor with three components: a saturation non-linearity, a phase-lead filter, and a threshold [23], giving rise to an output signal related to muscle force that is physiologically reasonable. In this study we include these to assess their impact on model computational performance, but they are not further used for feedback at this time, either to the model or to the user.

### B. Experiment 1: Kinematic fidelity of model-simulated hand motions

In the first experiment, the biomechanical model was driven with surface EMG signals recorded from normally-limbed individuals, and the simulated motions compared with those recorded from the participants using 3D motion analysis. EMG activity was recorded from four key muscles enabling the simulation of basic postures by the model. After giving informed consent for participation in the study (Keele Ethics reference ERP390), markers for motion capture and EMG recording electrodes were attached to the hand and forearm of participants (shown in Fig. 2).

**Fig. 2.**
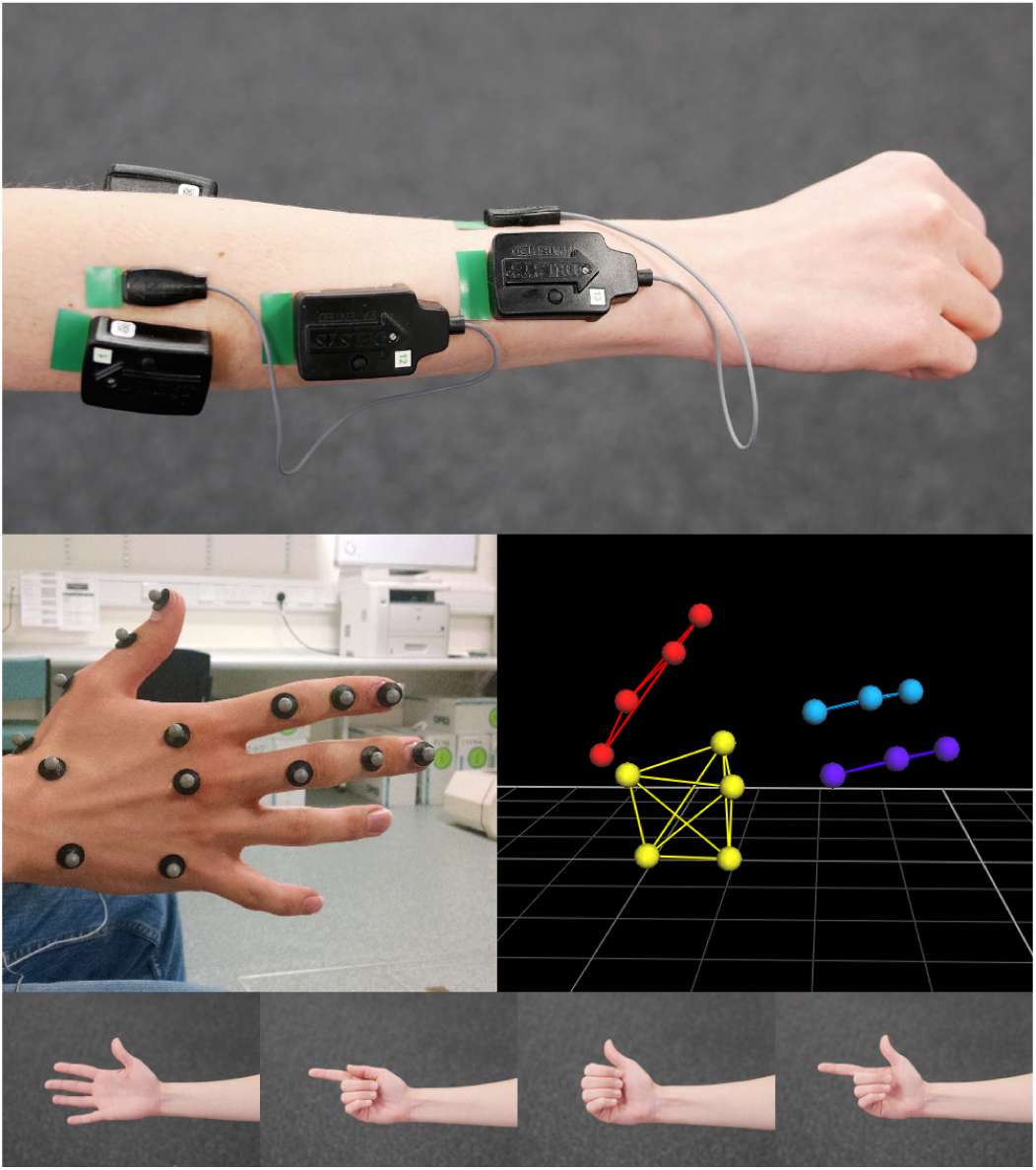
EMG electrodes were placed over four key muscle areas (top image) allowing independent control of the five postures (including rest). These were: the lateral part of EDC, the medial part of EDC, the FDS (just distal to the superficial wrist flexors), and the EPB. The middle row of images shows the locations of markers for the kinematic analysis, and the bottom row shows the four target postures presented.

For each participant, a static posture was recorded, with all digits extended. They were then asked to repeatedly open and close their hand for 30 seconds. Finally, they were asked to copy the hand movements presented to them in a demonstration video. The postures were presented to them in a randomised order over the course of 60 seconds, returning to a loosely closed posture in between. Thirty postures in total were attempted. The postures chosen were the fully open hand, pointing with the index finger, thumbs up, and an L-shape (pointing and thumbs up together). The resting posture was a loosely closed hand; the other postures are shown in the bottom row in Fig. 2. These postures were chosen as they were easy to achieve with the activation of superficial muscles that were feasible for recording via surface EMG. They are not intended to be a functionally comprehensive set, simply a feasible subset that demonstrate the performance of the real-time model as a controller for robotic hand movement.

We recorded the motions of the participants’ hands with a 3D motion analysis system (6-camera Vicon Bonita, Oxford Metrics Ltd), to compare the simulated motions with the recorded motions. Retroreflective markers of 6*mm* diameter were attached to the posterior surface of the palm, and the dorsal surface of the phalanges of the thumb, index and middle fingers. We did not include markers on the fourth and fifth digits, as they were not needed to identify the postures and were frequently occluded during the movements. This was a reduced version of the marker set described by Metcalf *et al.* [24].

The Datalink EMG system (Biometrics Ltd, Newport, Wales) was used to record muscle activity, and bipolar surface electrodes were placed over the following four muscles and muscle areas: extensor digitorum communis (EDC) lateral, approximately covering third, fourth and fifth digit; extensor digitorum communis (EDC) medial, capturing extension of the index finger; flexor digitorum superficialis (FDS) just distal to the wrist flexor bellies, capturing flexion of the fingers; extensor pollicis brevis (EPB), capturing extension of the thumb. Again, this is a reduced but feasible set of muscles that can be recorded independently with surface EMG electrodes, allowing us to demonstrate the performance of the model.

The EMG signals were rectified and processed with the use of a moving average filter with a window of 150*ms* (described in Blana *et al.* [25]). EMG amplitudes were scaled to a maximum contraction recorded for each posture to estimate normalised muscle activation from 0 to 1. The normalised EMG signal was then mapped to a combination of representative muscle tendon units (MTU) and these were used as inputs to the model; the outputs were the set of joint angles for all five digits. These mappings were selected to best achieve the desired movement with the limited surface EMG recordings, while keeping the control as intuitive as possible. The EMG to MTU mapping is shown in Table I.

**TABLE I.** Mapping of recorded EMG signals to model MTUs

| Source EMG | Modelled MTU |
| --- | --- |
| Extensor Digitorum Communis (lateral) | EDC (digits 3-5) |
| Extensor Digitorum Communis (medial) | EDC (digit 2) & EI |
| Flexor Digitorum Superficialis | FDS (digits 2-5) |
| Extensor Pollicis Brevis | EPB & EPL |

The trials with the static posture were used to scale the hand model to fit each participant with the OpenSim scaling method described in Delp *et al.* [19]). The dynamic trials were then used as inputs to inverse kinematic simulations with the scaled Opensim model. The outputs were joint angles.

As prosthetic hands typically have a single flexion/extension degree of freedom for each digit, we used a single joint angle to estimate the open/close movement of each digit: the thumb CMC joint, index finger MCP joint and middle finger MCP joint angles. The range of the three angles was found from the repeated open/close trial, and we used these to normalise the angles in the randomised posture trials, from zero (digit fully open) to one (digit fully closed).

Processed EMG data from the dynamic trials were input to the biomechanical model. The processing of the simulated angles was the same as the angles calculated from the Vicon recordings: the thumb CMC, index finger MCP and middle finger MCP model joint angles were normalised between zero and one using the range of simulated angles from the open/close trial.

We compared the recorded and simulated (normalised) movements throughout the randomised posture trials using the number of postures successfully achieved by each participant that were accurately replicated by the model. To do this, we assumed that if a (normalized) angle was below 0.2 the digit was open, if above 0.8 it was closed, and a particular posture was achieved if held for more than 0.2*s*. This allows for some variation in individual joint angles for the same posture that naturally occurs from trial to trial and between subjects, but is close enough to identify the posture. We then quantified the success rate as the ratio of the postures successfully adopted by the model to the postures attempted by the participant. Where the participant did not adopt the correct posture (or this was not clear from the Vicon data), we excluded that trial from the analysis. In addition to this, we report Pearson’s Correlation Coefficients for all movements, regardless of whether the posture was achieved by participant or model, to give an indication of the similarity of the movements made from one posture to another.

### C. Experiment 2: Real-time control of a robotic hand

In the second experiment, EMG recorded from a single, normally-limbed participant was used to drive a desktop robotic hand, via the biomechanical model, to assess the use of the real-time model with both robotic hardware and the user in the loop.

Surface EMG signals were recorded using a Trigno EMG system (Delsys Inc., Natick, MA) from the same four muscles used in Experiment 1. Processed EMG signals were used to drive a forward-dynamic simulation of hand dynamics, and the kinematic output from the model was then passed to the robotic hand so that the device mimicked the movements of the user. The robotic hand posture was updated every 100*ms*. The same set of movements was recorded as those used in Experiment 1; the EMG recording and processing were also the same, except for the addition of a minimum EMG threshold to remove low-level EMG activity. Low-level EMG fluctuations generate small forces in the model that cause the robotic hand to quiver: normalised EMG values below 0.05 were therefore set to zero to prevent this.

The robotic hand used was the Prensilia IH2 Azzurra. The hand consists of 5 degrees of freedom (thumb flexion/extension, thumb rotation, and flexion/extension of the second, third and combined fourth and fifth digits). It is cable actuated and features a built-in position controller that receives input via serial commands. The participant was shown a series of target postures by the experimenter, and asked to reproduce these with the robotic hand. The participant was prevented from seeing the robotic hand so that visual feedback did not influence the muscle activation patterns produced; the robotic hand was driven by the natural muscle activation patterns produced as a result of copying the experiementer’s target hand postures.

The performance of the model and hardware setup was quantified in terms of the time taken for each step in the data acquisition and processing cycle. The fidelity of reproduced movements were not quantified in this phase of the study; see Section II-B for those metrics.

### D. Experiment 3: Control of simulated gripping

In the final experiment, we created a simulation of object gripping, and explored the model’s use as a controller of grip force. In this simulation, a virtual (smooth-sided) cup was placed in the modelled hand, and slowly filled with water. A controller was included in the loop, and used to maintain grip force at a sufficient level to prevent slipping. The controller adjusted grip force by modulating the input muscle excitation level to the model during a forward-dynamic simulation (Fig. 3).

**Fig. 3.**
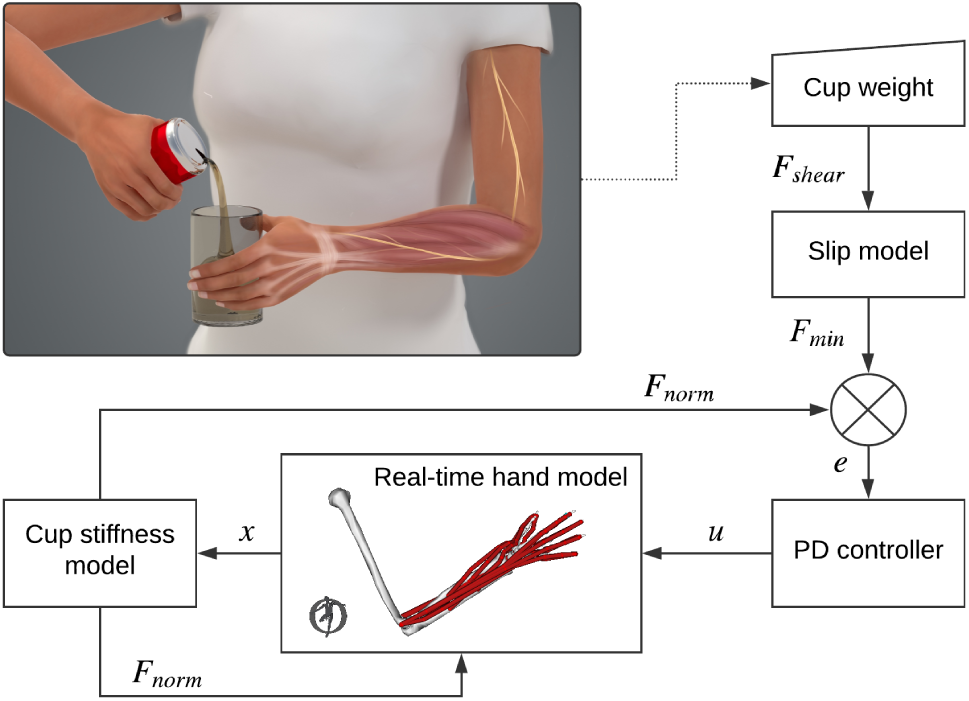
Schematic of simulated object gripping. The weight of the cup is altered to simulate filling, and the resultant shear force, *F*_*shear*_, on the finger is used to estimate the minimum contact force, *F*_*min*_, necessary to prevent slip. Muscle excitations, *u*, are input to the model and the resultant finger displacement, *x*, is output, from where cup stiffness is used to estimate fingertip contact force, *F_norm_*. A PID controller modulates the muscle excitation to ensure the contact force is kept above the minimum.

The presence of the object (a cup) was simulated by applying a force to the tips of the fingers in the model. A linear stiffness was assumed for the cup, allowing fingertip forces to be calculated from their displacement as the cup was squeezed. Fingertip forces were fed back to the model and the grip force modulated by controlling muscle excitations of the deep flexor muscles. Grip was maintained with the minimum force necessary to prevent slipping of the cup. The normal force required to achieve this was calculated by monitoring the weight of the cup and hence the shear (friction) force on the fingers, using an assumed value of *µ* = 0.3 for the coefficient of friction. The cup had a stiffness of 10*KN/m*, and the weight was varied to simulate filling with water.

The weight of the cup was evenly distributed between the four fingers on one side and the thumb on the other. The four fingers thus equally shared 50% of the weight. We focus on the index finger here for illustration, but the simulation involved all the fingers. The neural excitation of the Flexor Digitorum Profundus Indicis (FDPI) muscle was continually updated by a PD controller with proportional gain *K*_*p*_ = 0.01*N* ^-1^ and derivative gain *K*_*d*_ = 0.0005*N* ^-1^. The controller was tuned to give a short rise time and minimal oscillation.

The results of this experiment were assessed in terms of the computational performance of the model. In order to acheive real-time performance, the maximum stable step size for the forward-dynamic simulation needs to be larger than the amount of time needed to complete the computation of system dynamics. In addition, qualitative assessment of the grip force modulation behaviour in relation to normal grip was undertaken.

## III. Results

### A. Kinematic fidelity of model-simulated hand motions

A convenience sample of six normally-limbed individuals with no history of injury to the measured limb were recruited to the study to evaluate the kinematic fidelity of modelled hand motions. The mean age of the participants was 29.0(±6.2) years; three were male, three were female.

Fig. 4 shows the muscle excitation signals that are used as inputs for the model plotted over the raw EMG signals recorded from the corresponding muscles. The segment shown is for the ‘thumbs up’, ‘hand open’, ‘pointing with index finger’ and ‘L-shape’ postures.

**Fig. 4.**
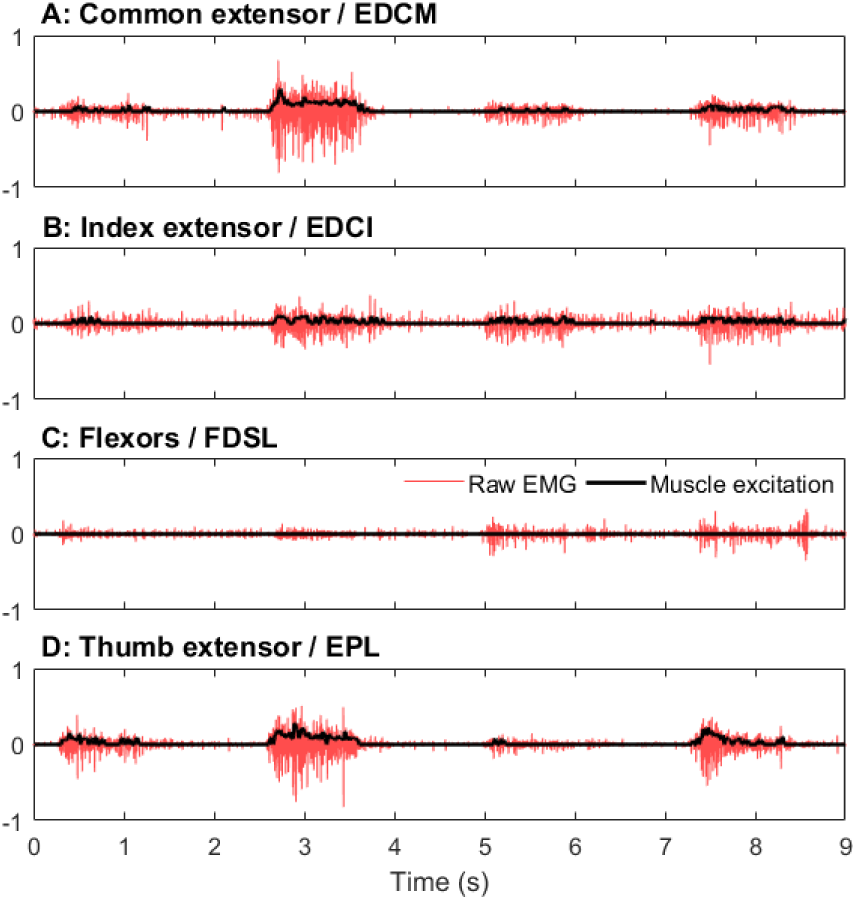
Example of the muscle excitation signals used as model inputs, together with the raw EMG signals from which they are derived. The segment shown includes the ‘thumbs up’, ‘hand open’, ‘pointing with index finger’ and ‘L-shape’ postures.

Fig. 5 shows a comparison of the measured hand kinematics against those estimated by the model for the same sequence of movements. The hand angles are normalised to the minimum and maximum values encountered during full opening and closing of the hand.

**Fig. 5.**
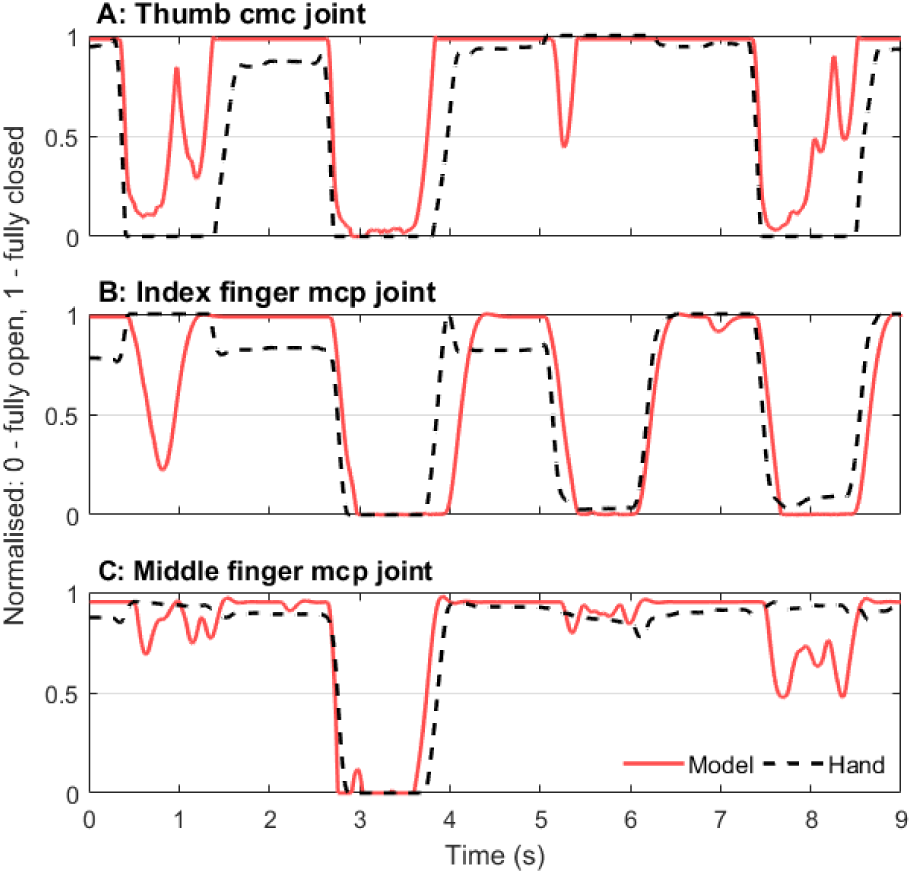
The normalised angles for both the measured hand kinematics and the model-predicted joint postures. These are for the same segment of data as shown in Fig. 4, normalized using the range estimated from the repeated open-close trial.

The success rate for the model matching the postures achieved by the participants is shown in Table II. In a few cases, the participant was not able to make all 30 target postures presented to them; the failed postures have been excluded from the analysis as the model posture could not be evaluated in those cases. Pearson’s Correlation Coefficient between the recorded and model-estimated angles for all movements was also used to estimate the fidelity of model-predicted movements; these are shown in Table III.

**TABLE II.**
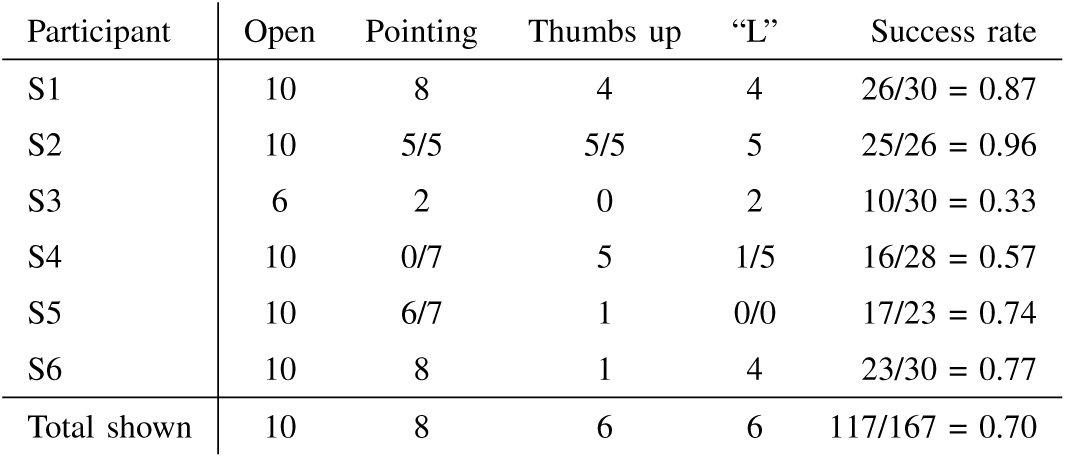
The success rate of the simulated posture matching the recorded posture. In some cases, the participant failed to achieve the target posture; those cases were excluded from the analysis.

| Participant | Open | Pointing | Thumbs up | “L” | Success rate |
| --- | --- | --- | --- | --- | --- |
| S1 | 10 | 8 | 4 | 4 | 26/30 = 0.87 |
| S2 | 10 | 5/5 | 5/5 | 5 | 25/26 = 0.96 |
| S3 | 6 | 2 | 0 | 2 | 10/30 = 0.33 |
| S4 | 10 | 0/7 | 5 | 1/5 | 16/28 = 0.57 |
| S5 | 10 | 6/7 | 1 | 0/0 | 17/23 = 0.74 |
| S6 | 10 | 8 | 1 | 4 | 23/30 = 0.77 |
| Total shown | 10 | 8 | 6 | 6 | 117/167 = 0.70 |

**TABLE III.** Pearson’s Correlation Coefficient for the model-predicted angles for each subject and each digit across all movements

| Participant | Thumb | Index | Middle |
| --- | --- | --- | --- |
| S1 | 0.96 | 0.95 | 0.99 |
| S2 | 0.96 | 0.96 | 0.99 |
| S3 | 0.89 | 0.98 | 0.93 |
| S4 | 0.93 | 0.92 | 0.98 |
| S5 | 0.89 | 0.97 | 0.99 |
| S6 | 0.96 | 0.91 | 0.99 |
| mean±std | 0.93±0.03 | 0.95±0.03 | 0.98±0.02 |

Finally, the addition of proprioceptive feedback in the form of the muscle spindle model allowed us to estimate the spindle firing rates from both primary and secondary afferents associated with these movements. Although they were not further used in this study, they are shown in Fig. 6 for reference.

**Fig. 6.**
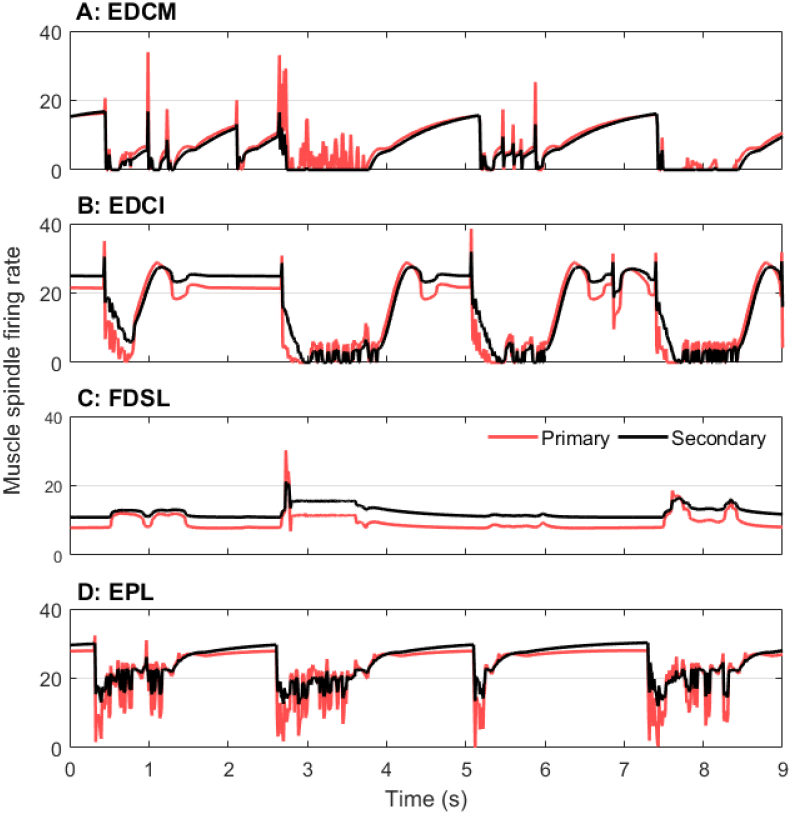
The model also simulates proprioceptive feedback. This shows the muscle spindle primary and secondary afferent firing rates for the same segment of data.

### B. Real-time control of a robotic hand

In the second experiment, EMG signals recorded in the same way as in Section II-B were again used to drive the forward-dynamic model, and the kinematic outputs from the model were passed to the robotic hand (Prensilia IH2 Azzura). This allowed the participant to directly control the movements of a physical, robotic hand in real time using forearm EMG signals. A small time delay associated with the EMG-envelope calculation was observed, but otherwise robotic hand movements mimicked those of the natural hand.

Fig. 7 shows the sequence of movements made by the user in controlling the hand, together with the actual posture adopted by the robotic hand. A full video of this is available in the Supplementary Material.

**Fig. 7.**
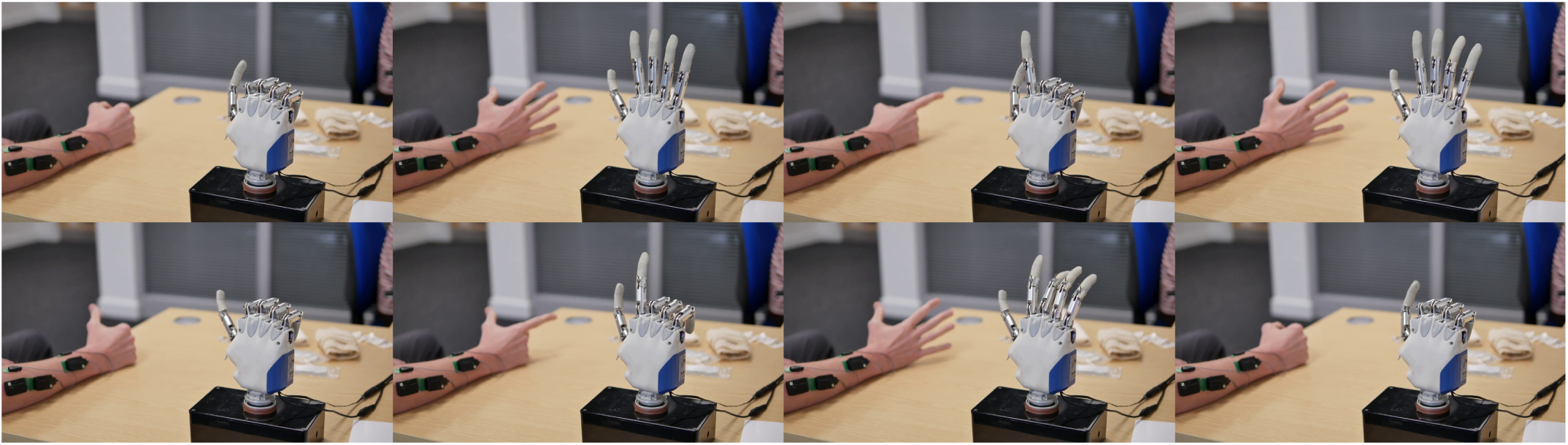
The Prensilia IH2 Azzurra robotic hand and the participant’s hand, shown in the various postures encountered during the trial. A video of the control achieved using this hand is available in the Supplementary Material.

Table IV shows the time required for each prosthesis control component, including the EMG acquisition and processing, dynamic simulations with the biomechanical hand model, and updating the robotic hand position. Out of 100*ms*, these processes take 16.2*ms* in total. The calculation of the pro-prioceptive feedback takes 2% of the time of the dynamic simulations.

**TABLE IV.** The computation time required for specific tasks in the control of robotic hand hardware.

| Process | Execution time (ms) |
| --- | --- |
| EMG acquisition and processing | 1.6 |
| Simulation of hand dynamics (25 timesteps) | 12.3 |
| Updating robotic hand position | 2.3 |
| Total | 16.2 |

### C. Control of simulated gripping

Fig. 8 shows the results of simulated gripping, where the amount of liquid in the cup is steadily increased. The initial value of 2*N* is the weight of the empty cup; the weight increases as the cup fills (Panel A). Panel B shows the activation of the FDPI muscle resulting from the changing force feedback. The initial spike is a response to the applied step load when the weight of the cup is placed in the hand. There is then a slow rise in activation as the cup fills. Panel C shows the fingertip force (solid line) maintained to just exceed the minimum force necessary to prevent slip (dashed line). Panel D shows the force in the FDPI tendon and Panel E the simulated output from the Golgi Tendon Organ Model.

**Fig. 8.**
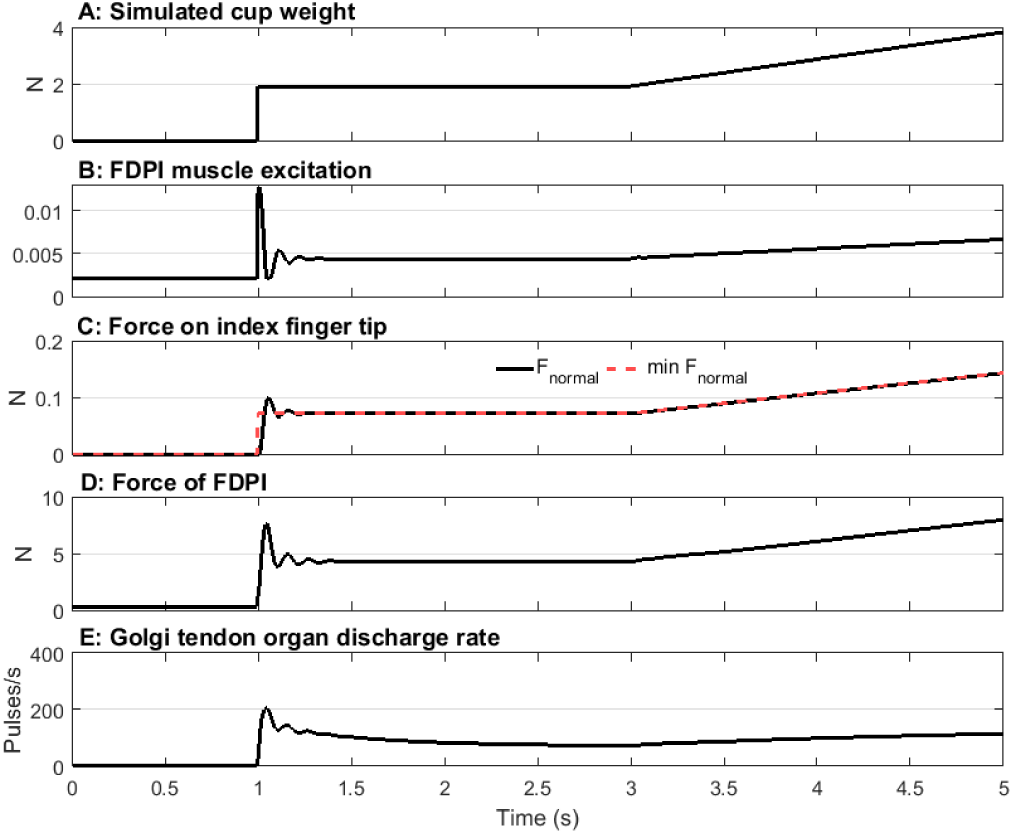
Panel A shows weight of the cup as liquid is added. Panel B shows activation in the FDPI muscle to ensure that the fingertip force just exceeds the minimum necessary to prevent slipping. Panel C shows the resulting fingertip force, Panel D the force in the finger flexor and Panel E the simulated GTO output.

Table V shows the computational performance achieved during the simulation of grip force control. The gripping task includes the simulation of the deformation of the cup used to estimate the fingertip contact force. Note that in ultimate use, this would not be simulated but measured, hence the key value in this table is the time taken to simulate the hand dynamics. Since this is 3.3*ms* and the integration step size is 4*ms*, the simulation is fast enough to run in real time.

**TABLE V.** The computation time required for specific tasks in the simulation of grip force control.

| Process | Execution time (ms) |
| --- | --- |
| Simulation of hand dynamics | 3.34 |
| PD Control execution | 0.009 |
| Simulated gripping task | 3.22 |
| Total | 6.57 |

## IV. Discussion

The aim of this study was to extend our initial work on real-time biomechanical simulation of hand dynamics to answer key questions regarding the feasibility of this method for prosthesis control. We have conducted three experiments intended to demonstrate (i) the fidelity of EMG-driven model movements in reproducing actions produced by normally-limbed volunteers; (ii) the computational performance of the model and its suitability for robotic hand control with hardware and the user in the loop; and (iii) the potential of the model-based control method to produce controlled gripping of objects in the hand.

In the first experiment, the model-predicted movements matched the recorded movements made by the subjects with good fidelity (Pearson’s Correlation Coefficient greater than 0.89 for all subjects) when continuous movement is compared. When comparing the final postures of the subjects’ actions with those from the model, the agreement appears somewhat less good, with the overall success rate on average being 0.70, but dropping as low as 0.33 for one subject. In some cases, this can be explained by the final posture falling just short of the threshold for classification of a given movement, although the continuous angle comparison may indicate better performance. Although a correlation coefficient of 0.89 is considered good and might be expected to lead to a close match between actual and desired movements in practice, the effects of this on actual use of a prosthetic hand will need to be assessed in terms of the functional, rather than kinematic, performance in future work. Related, preliminary work from our group has shown that control is acheivable with correlations in excess of approximately 0.55.

Furthermore, for some subjects with smaller arms, it proved difficult to separate the EMG signals into distinct functional movements, and this is an unsurprising limitation of the approach using discrete surface EMG recordings. The longer-term goal of this work is to make use of recorded motor control signals with greater spatial resolution by means of implanted electrodes, or even nerve recordings, and in that case EMG cross talk will be less of an issue. In this experiment we have focussed on a deliberately limited subset of movements to demonstrate the potential of the model-based approach, acknowledging that not all functional movements can be completed in this way.

In the second experiment, model simulations were amply fast, and in fact would have allowed for a much faster control loop than the one we chose (100*ms*). Kinematic outputs produced by the model were transmitted to the robotic hand via serial communication and the postures adopted almost instantaneously due to the robotic hand’s in-built controller. The time taken to update the robotic hand’s position was very small, as was the time to read and process EMG data. As expected, most of the time was spent simulating musculoskeletal dynamics. However, even this was well below what was necessary, and allows for a significant increase in model complexity if required, or a significant reduction in the robotic hand update frequency. This may allow for more responsive behaviour in real-world use.

In the final experiment, we have demonstrated the potential of the model in modulating finger flexor force in a simulation of closed-loop control of gripping. This shows that the model could be used to regulate muscle activity in response to the changing demand for grip force. The inclusion of the muscle model, with its first order delays and elastic tendon, gives rise to a compliant, human like grip that may lead to more natural control. This demonstration features feedback-only control of grip, whereas human grip features much more influence from feedforward mechanisms, and recent work has shown the importance of both feedforward and feedback mechanisms for improving control of grip in upper limb prosthesis users [26]. In our long-term vision, descending commands recorded from the user would provide the feedforward element, and force-related signals from simulated GTO and haptic sensors in the fingertips of the prosthetic device would be fed back. This means the user themselves would modulate descending commands controlling muscle activation, and the need for the PD controller is removed. This brings the control of slip-prevention to the user, rather than leaving the prosthetic device itself to autonomously control grip.

The use of a biomechanical model driven by cognate muscle activity suggests that the movement of the robotic or prosthetic hand should match the naturally intended movement of the user, and indeed to some extent we have demonstrated that to be the case. However, it should be noted that no attempt at model customisation beyond simple scaling was made in this case, so some degree of learning or adaptation to the differences between the natural hand and the model may be expected. Since our goal is to enable prosthesis control in someone without a natural hand, this may be of secondary importance compared to the effect of learned non-use in that person. Indeed, supporting this, recent studies have shown that an amputee may have more difficulty in controlling coordinated movement than normally-limbed individuals [16], [27]. Nonetheless, scaling to the contralateral limb could be carried out to reduce initial differences in control signals and minimise the learning required.

Recently published work [17] has shown the effectiveness of a similar biomechanical model approach to prosthesis control in user testing with amputees, albeit focusing on whole hand movement and not individual finger control. In that study, the simulation stopped short of full rigid-body kinematics, but transferred joint torques directly to the prosthetic device by controlling joint velocity. Thus the need for numerical integration of the state dynamics was effectively replaced with a physical model in the form of the prosthesis. This is an elegant solution that allows robust, pragmatic control.

Our approach uses knowledge of limb biomechanics to provide control signals for a prosthetic device via a real-time model, and allows simulation of prosthesis function also in the absence of the physical device. This may allow user training and system optimisation to take place ahead of fitting, minimising the learning time for the user. In our long-term vision, recordings of descending commands made either from residual muscle and nerve, or muscles innervated through targeted reinnervation [13], will produce the driving signals for the model. Furthermore, we have shown the ability to include physiologically meaningful simulations of proprioceptive output of both joint kinematics (via muscle spindle output) and force feedback (GTO outputs) that could be fed back to the user by peripheral nerve stimulation in future work. This will allow truly closed-loop control of hand function for prosthesis users, enabling state-of-the-art devices to be used to their full potential.

## V. Conclusions and future work

In this study, we have demonstrated the feasibility of real-time biomechanical simulation of hand function for prosthesis control by showing good fidelity between model-predicted and human measured kinematics, faster than real time computational performance, and the potential for model-based grip-force control. A number of limitations of the model have been identified in terms of access to driving signals, but the potential for enabling greater dexterity as device-human interfacing improves is clear.

Future work will involve testing the functional performance of this approach in upper limb amputees both in simulation and with state-of-the-art prosthetic devices. Many questions remain regarding access to and user control of suitable signals, as well as users’ ability to learn the complex dynamics of multi-joint movement. We have also shown the potential to simulate meaningful feedback signals, and future work will need to assess the viability of delivering and interpreting these.

To facilitate shorter term translation of this work to current devices where invasive recordings may not be desirable or possible, further work investigating the improved extraction of information from surface EMG recordings should be pursued. This could include combining the musculoskeletal modelling approach with computational intelligence to infer information on deeper muscles that are not amenable to surface recordings.

## Supporting information

Video of robotic hand being controlled by user EMG signals

## Acknowledgment

The authors gratefully acknowledge the support of the Engineering and Physical Sciences Research Council (EP/M025977/1) and the National Institutes of Health (NIH-5R01EB011615) in this research.

